# Selection and validation of reference genes for quantitative real-time polymerase chain reaction in *Serratia ureilytica* DW2

**DOI:** 10.1101/2022.05.03.490547

**Authors:** Fenglin Bai, Bianxia Bai, Tingting Jin, Guiping Zhang, Jiahong Ren

## Abstract

*Serratia ureilytica* DW2 is a highly efficient phosphate-solubilizing bacterium isolated from *Codonopsis pilosula* rhizosphere soil that can promote the growth of *C. pilosula*. However, no validated reference genes from the genus *Serratia* for use in quantitative real-time polymerase chain reaction (RT–qPCR) normalization have been reported. To screen stable reference genes in *S. ureilytica* DW2, the expression of eight candidate reference genes (16S rRNA, *ftsZ, ftsA, mreB, recA, slyD, thiC*, and *zipA*) under different treatment conditions (pH, temperature, culture time, and salt content) was assayed by RT–qPCR. The expression stability of these genes was analyzed with different algorithms (geNorm, NormFinder, and BestKeeper). To verify the reliability of the data, the most stably expressed reference gene was used to quantify expression of the glucose dehydrogenase (*gdh*) gene under different soluble phosphate levels. The results showed that the *zipA* and 16S rRNA genes were the most stable reference genes, and the least stable were *thiC* and *recA*. The expression of *gdh* was consistent with the phosphate solubilization ability on plates containing National Botanical Research Institute phosphate (NBRIP) growth medium. Therefore, this study provides a stable and reliable reference gene for *Serratia*, which is vital for the accurate quantification of functional gene expression in future studies.

## Introduction

Real-time quantitative polymerase chain reaction (RT–qPCR) is an established technique for the quantitative expression analysis of target genes under different treatment conditions [1]. This technique has many advantages, including high sensitivity, accuracy, repeatability, and low labor intensity [2–6]. However, RT–qPCR can be affected by RNA quality, amplification efficiency, and the stability of reference gene expression among different samples [7–9]. Therefore, reliable reference genes are vital to accurately compare the gene expression level between control and experimental groups [10]. The ideal reference genes are usually stable under different experimental conditions, with abundances closely correlated with total mRNA amounts [7]. These genes, such as *gyrA, gyrB, recA, fabD, rpoB*, and 16S rRNA, are usually used as reference genes, but they are not universal for all bacteria [11]. As there have been no reports related to the reference genes for the genus *Serratia*, it is necessary to explore potential reference genes for this genus.

*Serratia ureilytica* DW2 is an excellent phosphate-solubilizing bacterium (PSB) previously isolated from the *Codonopsis pilosula* rhizosphere by this research group. In greenhouse experiments, the strain DW2 could promote the growth of *C. pilosula* after inoculation in rhizosphere soil (unpublished). *C. pilosula* is a traditional Chinese medicine that is used widely in clinical therapeutics to enhance immunity, lower blood pressure, and improve microcirculation[12]. *C. pilosula* is cultivated in large quantities to meet high market demand. During cultivation, fertilizer is often used to improve yield, which affects the quality of *C. pilosula* and its soil environment. *S. ureilytica* DW2 could be used to develop biofertilizers for *C. pilosula* and thereby promote its sustainable cultivation.

To further study the growth-promoting characteristics of *S. ureilytica* DW2 at the molecular level, reliable reference genes need to be confirmed. Reference genes can be used to validate the gene expression among different samples. Therefore, in the present study, eight genes, namely, recombinase A (*recA*), 16S ribosomal RNA (*16S rRNA*), rod shape-determining protein MreB (*mreB*), FKBP-type peptidyl-prolyl cis-trans isomerase (*slyD*), hydroxymethylpyrimidine phosphate synthase (*thiC*), cell division protein (*zipA*), cell division protein FtsA (*ftsA*), and FtsZ (*ftsZ*), were selected as candidate reference genes. Their expression was detected by RT–qPCR, and their expression stabilities were evaluated by three methods (geNorm, NormFinder, and BestKeeper). This identification of reference genes in *S. ureilytica* DW2 is the first reported research on reference genes in the genus *Serratia* and could provide a basis for the quantification of functional gene expression in this genus.

## Materials and methods

### Selection of reference genes and design of primers

Eight genes (16S rRNA, *ftsZ, ftsA, mreB, recA, slyD, thiC*, and *zipA*) were selected for the real-time RT–qPCR-based relative expression analysis of target genes [13–15]. The sequences of these 8 candidate reference genes were obtained from *S. ureilytica* DW2 genome data. Primers for candidate reference genes were designed using Primer 3 software and synthesized by Suzhou Jinweizhi Biotechnology Co., Ltd. (China).

### Bacterial strain and culture conditions

Sample preparation of candidate reference genes under different expression conditions. *S. ureilytica* DW2 was cultured in LB solid medium overnight, and single colonies were selected and activated in liquid LB at 30°C overnight 2 times. The cells were inoculated at 1% (volume fraction) in a triangular flask of 200 mL liquid LB, starting at pH 7, and cultured at 30°C for 9 h to logarithmic metaphase. At different temperatures (25, 30, and 37°C), NaCl concentrations (0.5%, 1.0%, and 1.5%), pH values (5, 7, and 9), and culture times (12, 24, and 36 h), single factor analysis was used for grouping. The samples were centrifuged at room temperature at 6 000 r/min for 10 min, frozen with liquid nitrogen and stored in a −80°C refrigerator for later use.

### RNA extraction and cDNA synthesis

Two milliliters of DW2 cells was collected by centrifugation. RNA extraction was performed as follows: total RNA was extracted using TRIzol reagent (Invitrogen, USA) according to the manufacturer’s instructions. To ensure the complete removal of any contaminating DNA, all RNA preparations were subjected to a second DNase treatment using the genomic DNA (gDNA) Eraser (Perfect Real Time Kit, Takara Biochemicals, China). RNA concentrations were quantified on a Nanodrop 2000 spectrophotometer (Thermo Fisher Scientific, USA). The OD260/OD280 and OD230/OD260 ratios were calculated to evaluate the purity of the RNA extracts. The integrity of the RNA was checked by agarose gel electrophoresis. Complementary DNA (cDNA) was synthesized using the PrimeScript RT reagent kit (Perfect Real Time Kit, Takara Biochemicals, China). According to the vendor’s instructions, 1000 ng total RNA was reverse transcribed using Phusion RT–PCR Kit (Finnzymes, Finland) random primers.

### RT–qPCR run conditions

RT–qPCR was performed on a 7500 Fast Real Timer PCR system (ABI, USA) using a SYBR® Premix Ex Taq™ kit (Takara, China). The standard curve was measured with serial 10-fold dilutions of cDNA to determine the associated amplification efficiency (E) and nonspecific products. The PCR consisted of 10 μL of SYBR Green PCR Master Mix, and the upstream and downstream primers were 0.5 μL each, 0.2 μL of ROX (passive reference dye), and 2.0 μL of diluted cDNA template in a total volume of 20 μL. The PCR amplification process was as follows: 95°C 5 min; 40 cycles of 95°C 5 s denaturation, 60°C 34 s annealing extension. All qRT–PCR products were analyzed in three technical replicates and validated by subsequent melting curve analysis.

### Analyses of reference gene expression stability

Three software programs (geNorm [16], NormFinder [17], and BestKeeper [18]) were used to compare the expression stability of eight candidate reference genes under 12 treatments.

geNorm uses expression stability measurement values (M) to evaluate candidate reference genes. The most stable expression is indicated by the lowest M value. The algorithm defines the stability of a gene by calculating the average paired variation, whose cutoff value is 1.54 [18]. When the critical value is greater than 1.5, the gene is not suitable for use as an internal reference. In the range of m values less than 1.5, the smaller the value is, the better the gene stability and the more suitable it is for use as an internal reference gene.

With NormFinder calculation, two stable internal parameters can be selected. Unlike geNorm, NormFinder takes into account changes in expression within and between groups. Therefore, the misinterpretation caused by gene coregulation can be avoided by this algorithm. Similar to geNorm, NormFinder analyzes candidate reference genes according to their expression stability value, and the higher the expression stability, the lower the value, and vice versa [17].

BestKeeper calculates the coefficient of variation (CV), Pearson correlation coefficient (r) value, geometric mean (GM), arithmetic mean (AM) and other rich data. Suitable reference genes should meet both the normalization factor and pairwise correlation analysis limitations. This method compares the standard deviation of each gene, takes a standard deviation (SD) less than 1 as a necessary condition and does not consider those with values greater than 1 as suitable internal references. On this basis, the gene with the minimum CV (%Cq) and ±Cq value was considered the most stable. BestKeeper analysis compared pairs, and the correlation coefficient (r) was calculated. The closer the r value was to 1, the more suitable the gene was as an internal reference.

### Phosphate-solubilizing ability under different concentrations of soluble phosphate

On NBRIP medium, the transparent circle formed by PBS bacteria can qualitatively represent the phosphorus solubilizing capacity of microorganisms, which is determined by the ratio of the transparent zone diameter and the colony diameter [19]. Ca_3_(PO_4_)_2_ was used as the only insoluble phosphate source in NBRIP solid medium. After adding soluble phosphate, K_2_HPO_4_, at three different concentrations (0, 5.0 and 10.0 mmol L^-1^), the transparent circles around colonies were observed after 7 days of culture to infer the phosphorus solubilizing ability of DW2. Three replicates were performed for each soluble phosphate level.

### Validation of glucose dehydrogenase (*gdh*) gene expression

PSB converts insoluble phosphate to soluble phosphate by producing organic acids. One of its pathways is the conversion of glucose to gluconic acid, catalyzed by glucose dehydrogenase (GDH). GDH is a membrane-binding enzyme that participates in the direct oxidative pathway of glucose catabolism. This study analyzed the genomic sequence of *S. ureilytica* DW2 and successfully cloned the *gdh* gene. According to the average Cq value of the reference genes obtained by screening, the relative expression level of *gdh* at different soluble phosphate levels was calculated by the 2^-ΔΔct^ method [20]. Biostatistics was used to analyze the significant difference in the relative expression levels of target genes under different levels of soluble phosphate. By comparing the qRT–PCR results of the target gene and the ratio of colony diameters, the expression stability of the reference genes obtained by screening was verified.

## Results

### Evaluation of experimental credibility

Information on the 8 candidate reference genes is shown in Table 1. The products of the 8 candidate reference genes were between 100 and 200 bp, which is consistent with the expected band size, and the bands were clear and single. (Fig. S1). Melting curve analysis showed that all primers were specific, as a single signal peak was obtained. The melting temperatures of the amplicons were in the range of 85.56– 90.17°C (Fig. S2). Standard curves showed that PCR efficiencies (E[%]) ranged from 91.398% (*thiC*) to 100.214% (*recA*). The correlation coefficient (R^2^) results ranged from 0.994 to 0.999, indicating that the data were reliable.

**Table 1.**
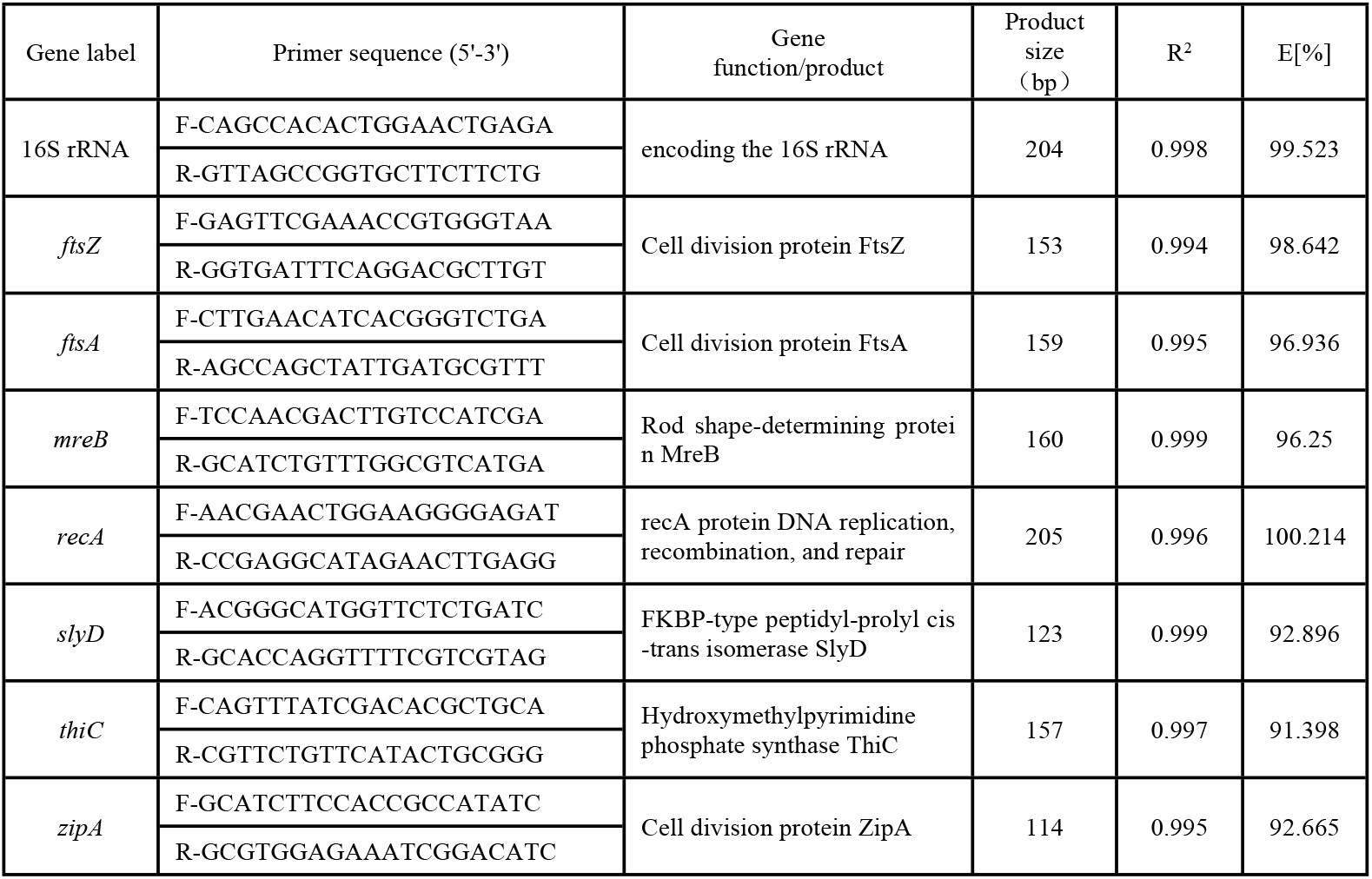
Primer list used for qPCR validation.

### Transcript levels of candidate reference genes

The expression levels of candidate genes in *S. ureilytica* DW2 were assessed by RT– qPCR. The expression of each candidate gene in the 12 processed samples is shown in Fig. 1. All genes were amplified prior to the amplification cycle of the PCR amplification profile. Among the eight candidate reference genes, the highest geometric mean of the Cq value was for *thiC* (32.57), and the lowest was for 16S rRNA (12.37). The Ct values of 16S rRNA ranged from 10.91 to 13.66, indicating higher expression levels than those of the other seven genes. Then, the data of 8 candidate reference genes were further analyzed.

**Fig. 1.**
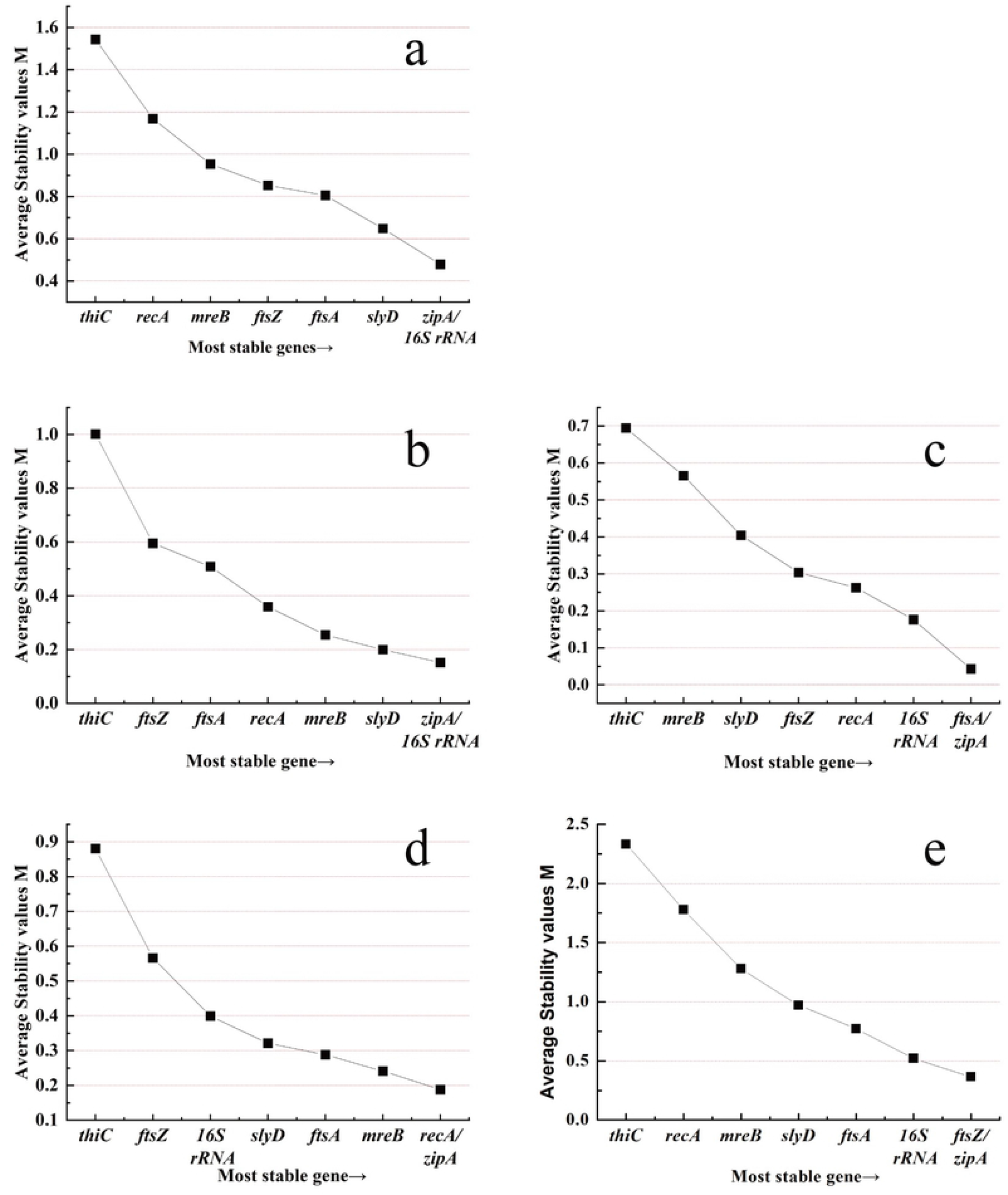
Quantification cycle (Cq) values of eight candidate reference genes.

### Comparison of the expression stability of candidate reference genes according to normalization strategies

Three independent statistical algorithms were used to compare the expression stability. geNorm is an effective tool for selecting stable internal parameters. It selects two targets with the same expression ratio as the most stable pair in the test samples [21]. Figure 2 shows the M values of the 8 reference genes in the primary selection. The M values of seven candidate genes were < 1.5, which indicates that they can theoretically be used as reference genes, with the exception of *thiC* (M 1.54). Overall, the least stable genes were *thiC* and *recA*. The 16S rRNA and *zipA* genes showed the lowest M values (0.48). Therefore, 16S rRNA and *zipA* were considered the two most stable reference genes according to the geNorm calculation.

**Fig. 2.**
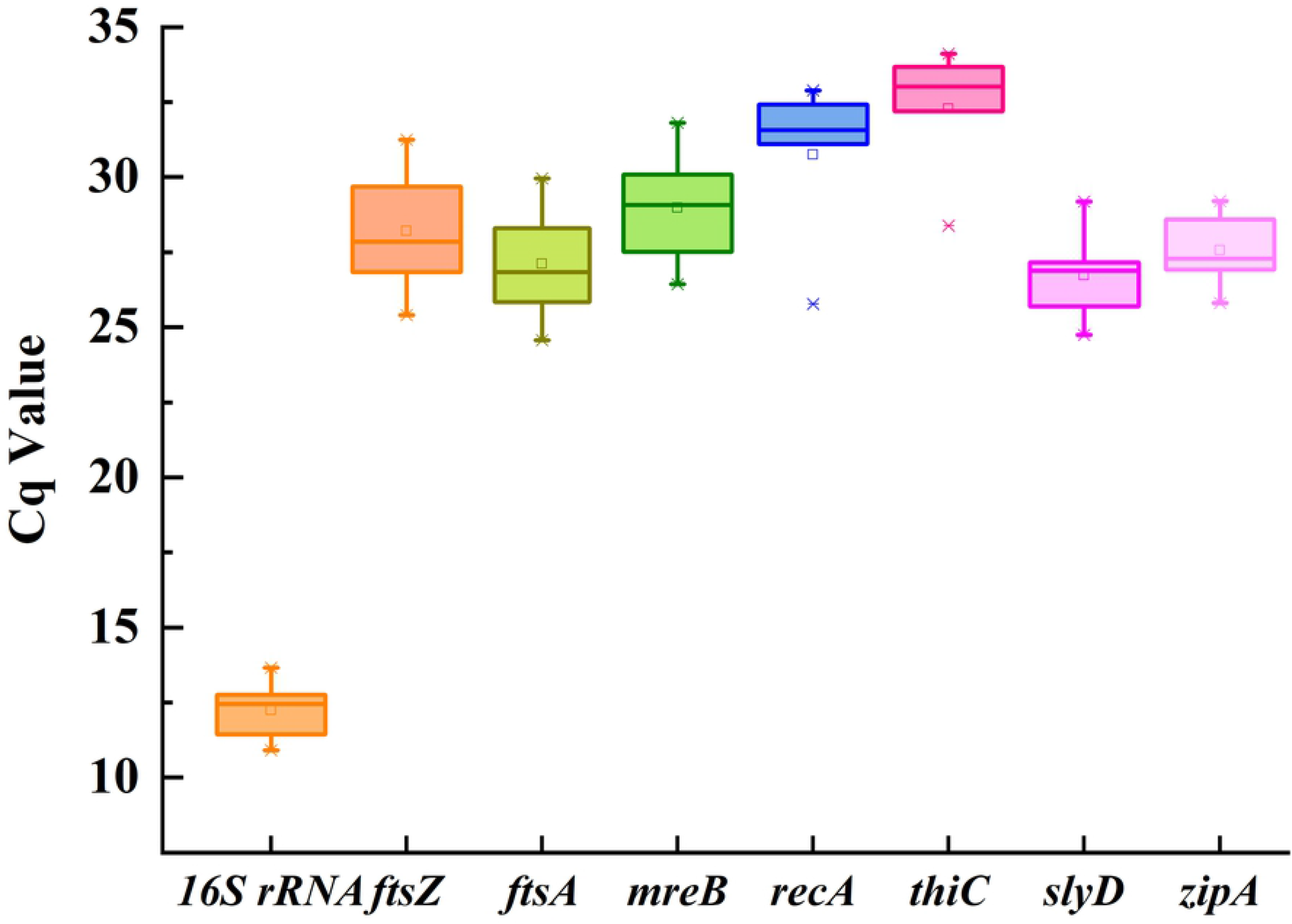
geNorm analysis of reference genes in *S. ureilytica* DW2. Analyses were conducted with geNorm software. A low stability M value (y-axis) indicates that the gene was stably expressed under the 12 experimental conditions.

NormFinder was also used to analyze the expression stability of the reference genes. Generally, the smaller the S value is, the higher the expression stability of the reference genes and the higher the rank [22]. According to NormFinder analysis (Table 2), the top three genes with the lowest S values were *slyD* (S = 0.106), *zipA* (S = 0.128), and *mreB* (S = 0.155) under all conditions. By comparison, *thiC* (S = 0.290) and *recA* (S = 0.277) had higher S values and ranked as the least stable.

**Table 2.** Stability values (S) of the eight candidate reference genes in *S. ureilytica* DW2 determined with NormFinder.

| Rank | Overall |  | pH values |  | Salt |  | Temperatures |  | Culture time |  |
| --- | --- | --- | --- | --- | --- | --- | --- | --- | --- | --- |
|  | Gene | Stability Value | Gene | Stability Value | Gene | Stability Value | Gene | Stability Value | Gene | Stability Value |
| 1 | *slyD* | 0.106 | *slyD* | 0.009 | 16S rRNA | 0.009 | *ftsZ* | 0.009 | *slyD* | 0.052 |
| 2 | *zipA* | 0.128 | *ftsZ* | 0.009 | *zipA* | 0.009 | *zipA* | 0.011 | *zipA* | 0.104 |
| 3 | *mreB* | 0.155 | *mreB* | 0.012 | *ftsA* | 0.010 | *recA* | 0.033 | *ftsZ* | 0.258 |
| 4 | *ftsZ* | 0.170 | *ftsA* | 0.015 | *ftsZ* | 0.030 | *ftsA* | 0.041 | *mreB* | 0.324 |
| 5 | 16S rRNA | 0.241 | *zipA* | 0.059 | *thiC* | 0.037 | *mreB* | 0.049 | 16S rRNA | 0.389 |
| 6 | *ftsA* | 0.251 | *recA* | 0.086 | *slyD* | 0.043 | *slyD* | 0.049 | *recA* | 0.393 |
| 7 | *recA* | 0.277 | *thiC* | 0.097 | *recA* | 0.043 | 16S rRNA | 0.063 | *thiC* | 0.405 |
| 8 | *thiC* | 0.290 | 16S rRNA | 0.170 | *mreB* | 0.099 | *thiC* | 0.335 | *ftsA* | 0.553 |

Comparing the *operation* results of geNorm and NormFinder under different treatment conditions (Table 2, Fig. 2), according to the geNorm algorithm, 16S and *zipA* could be considered reference genes under the different pH values. However, NormFinder advised against this, ranking 16S and *zipA* at positions eight and five, respectively. After salt treatment, *zipA* and *ftsA* showed the least overall variation under both algorithms. At different temperatures, *ftsZ* and *zipA* were the most stable under the two algorithms. The genes *slyD* and *zipA* performed the best at different culture times under both algorithms.

BestKeeper is a calculation method that mainly performs paired correlation and regression analyses on the selected genes [23]. The results of BestKeeper analysis are shown in Table 3. In general, ± Cq values for appropriate reference genes should < 1. In terms of this criterion, the genes 16S rRNA (0.8) and *zipA* (0.99) passed this screening. However, *ftsZ* has the highest SD [± Cq] (1.49). Moreover, the ± x-fold value should be < 2 [24], and only the 16S rRNA and *zipA* genes met this requirement. For the 16S rRNA and *zipA* genes, the CV (%Cq) values (3.60) of *zipA* were lower. A comparison of the r values of the reference genes shows that the r value of *zipA* (0.95) was closer to 1. The *thiC* gene was the least stable, with an r value of only 0.68. By BestKeeper’s calculation, *zipA* and 16S rRNA were considered the two best internal reference genes.

**Table 3.**
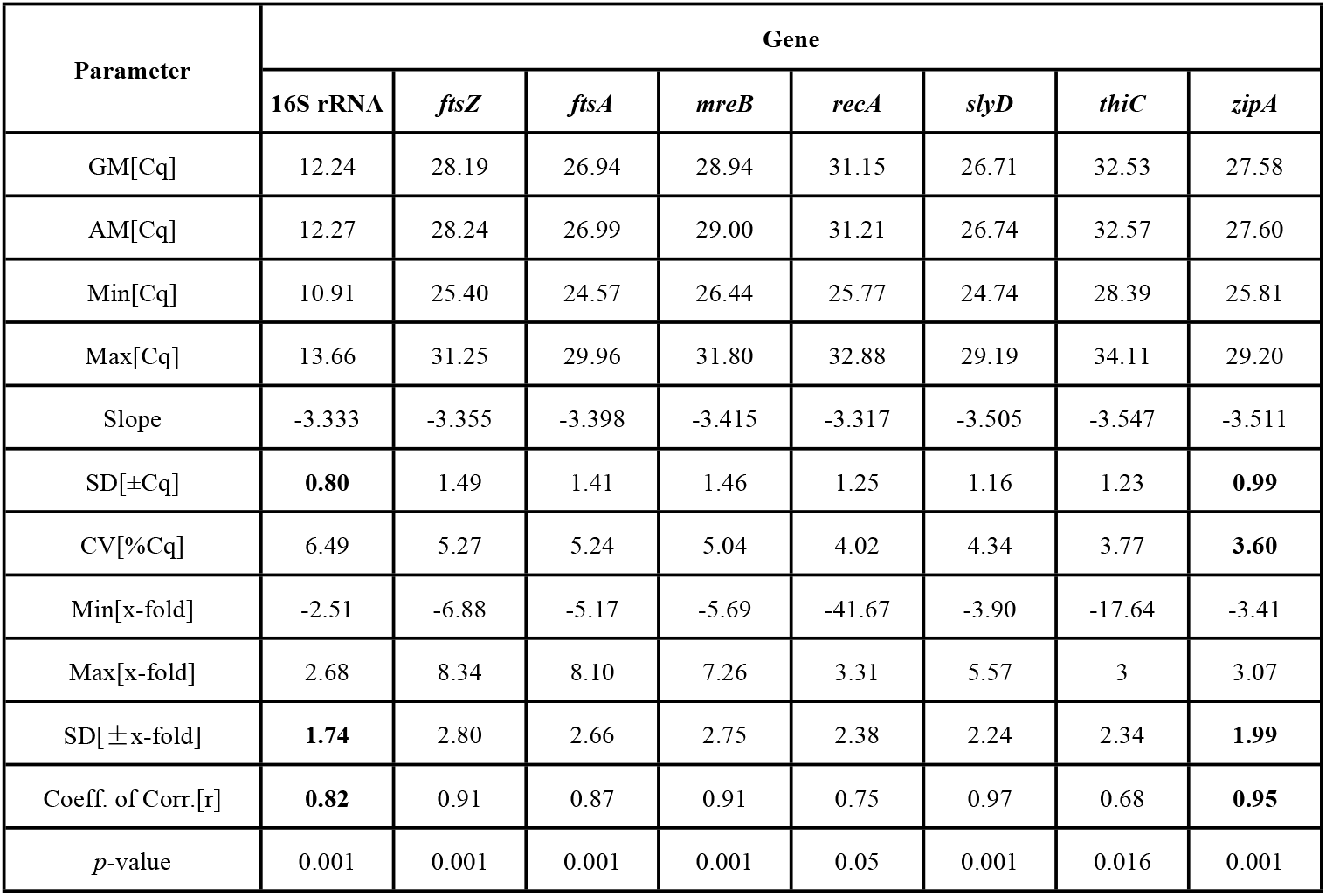
Descriptive statistics of candidate reference genes.

| Parameter | Gene |  |  |  |  |  |  |  |
| --- | --- | --- | --- | --- | --- | --- | --- | --- |
|  | 16S rRNA | *ftsZ* | *ftsA* | *mreB* | *recA* | *slyD* | *thiC* | *zipA* |
| GM[Cq] | 12.24 | 28.19 | 26.94 | 28.94 | 31.15 | 26.71 | 32.53 | 27.58 |
| AM[Cq] | 12.27 | 28.24 | 26.99 | 29.00 | 31.21 | 26.74 | 32.57 | 27.60 |
| Min[Cq] | 10.91 | 25.40 | 24.57 | 26.44 | 25.77 | 24.74 | 28.39 | 25.81 |
| Max[Cq] | 13.66 | 31.25 | 29.96 | 31.80 | 32.88 | 29.19 | 34.11 | 29.20 |
| Slope | -3.333 | -3.355 | -3.398 | -3.415 | -3.317 | -3.505 | -3.547 | -3.511 |
| SD[±Cq] | **0.80** | 1.49 | 1.41 | 1.46 | 1.25 | 1.16 | 1.23 | **0.99** |
| CV[%Cq] | 6.49 | 5.27 | 5.24 | 5.04 | 4.02 | 4.34 | 3.77 | **3.60** |
| Min[x-fold] | -2.51 | -6.88 | -5.17 | -5.69 | -41.67 | -3.90 | -17.64 | -3.41 |
| Max[x-fold] | 2.68 | 8.34 | 8.10 | 7.26 | 3.31 | 5.57 | 3 | 3.07 |
| SD[±x-fold] | **1.74** | 2.80 | 2.66 | 2.75 | 2.38 | 2.24 | 2.34 | **1.99** |
| Coeff. of Corr.[r] | **0.82** | 0.91 | 0.87 | 0.91 | 0.75 | 0.97 | 0.68 | **0.95** |
| *p*-value | 0.001 | 0.001 | 0.001 | 0.001 | 0.05 | 0.001 | 0.016 | 0.001 |

The pairwise variation (V value) method was used to calculate the optimal number of reference genes by geNorm. According to Vandesompele, no additional reference gene is required if Vn/Vn+1 is < 0.15 [16]. All samples were included to calculate the optimal number of reference genes (Fig. 3). Furthermore, geNorm indicated the recommended number of reference genes under various treatment conditions.

**Fig. 3.**
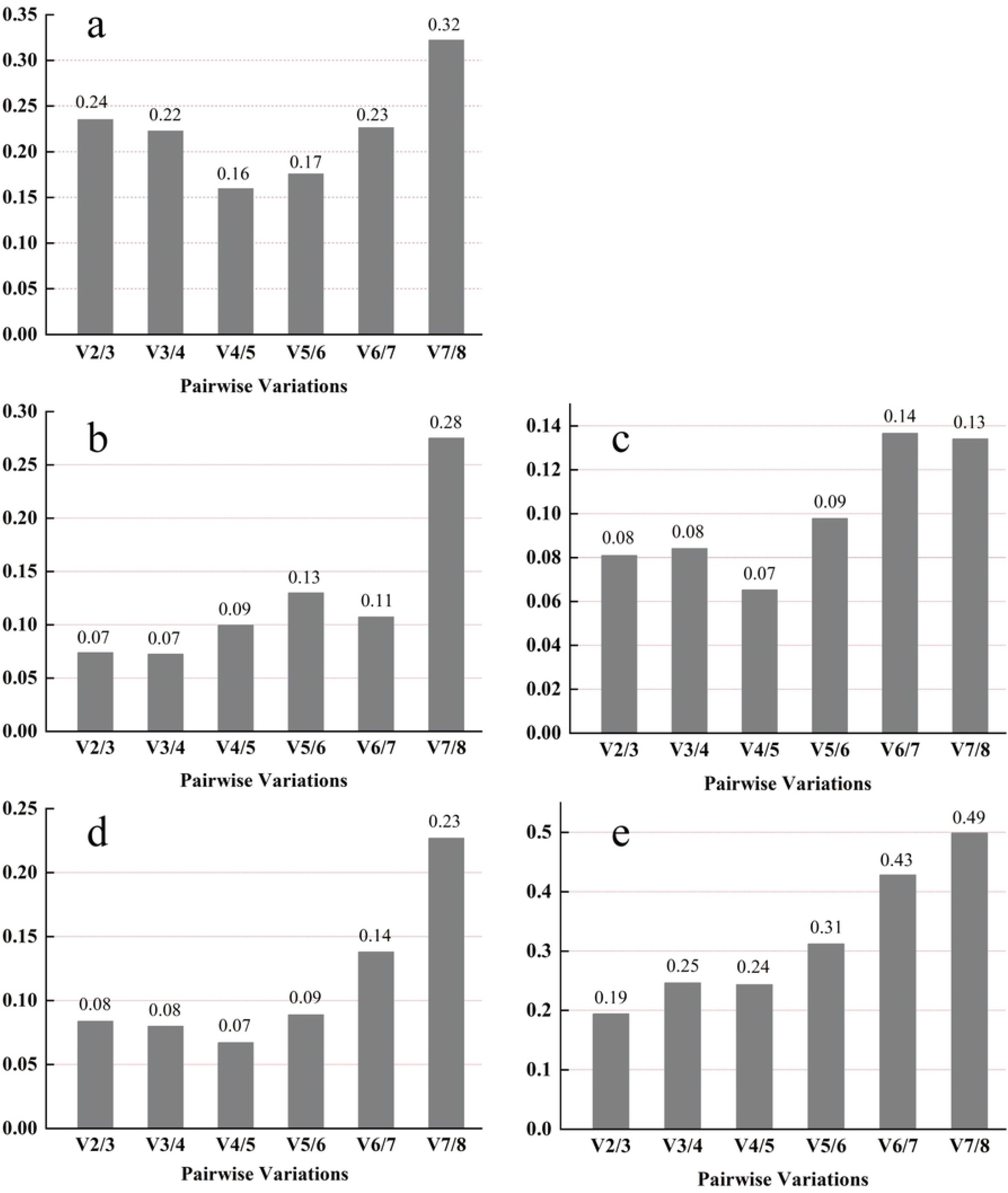
The pairwise variation method was used to calculate the optimal number of reference genes. Under all conditions (a). Treatments with temperatures (b), NaCl concentrations (c), pH values (d), culture times (e).

Under the 12 treatment conditions, the V2/V3 value was much lower than the cutoff value, which suggests that the first two genes were sufficient for use as internal references. When analyzing all the different conditions, the values of Vn/Vn+1 ranged from 0.16 for V4/5 to 0.32 for V7/8. The value for V4/V5 (0.16) was higher than the cutoff of 0.15. However, the recommended V value (0.15) is only a recommended critical point and should not be used as a strict cutoff value [16].

According to the geometric mean of the ranking obtained by these three programs (Table 4), *zipA* and 16S rRNA had the smallest geometric mean and the highest expression stability.

**Table 4.**
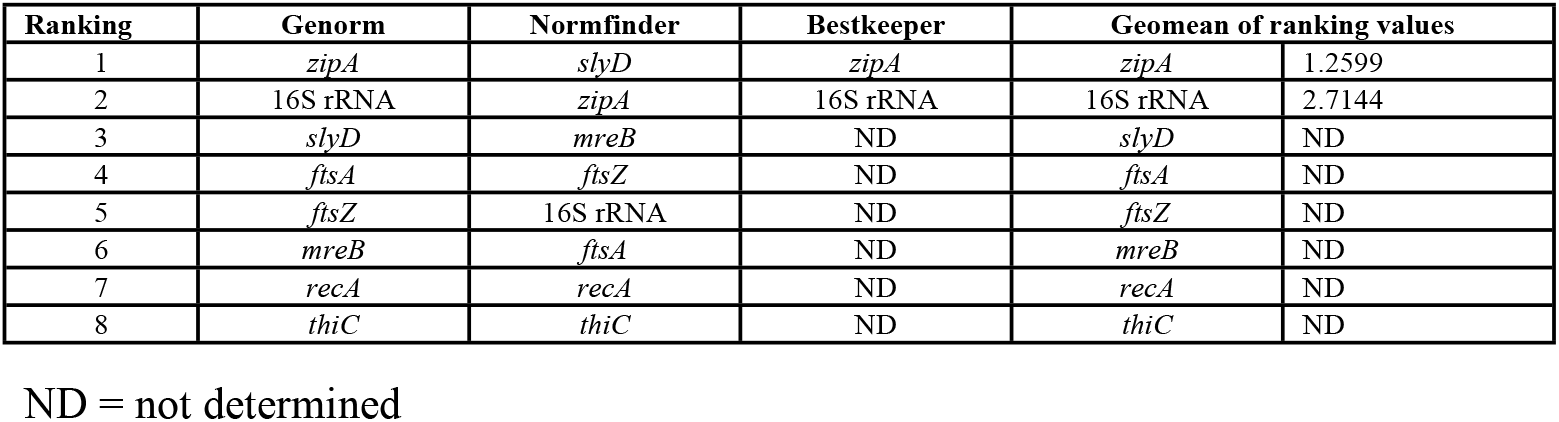
Ranking of the candidate genes based on the results of analyses with geNorm, NormFinder and BestKeeper.

### Validation of *gdh* gene variation under different soluble phosphate levels

Phosphate-solubilizing bacteria can dissolve the inorganic phosphorus fraction into available phosphorus for plants and thereby indirectly promote plant growth [25]. The phosphate-solubilizing activity of phosphate-solubilizing bacteria is affected by the amount of exogenous soluble phosphate.

*zipA* and 16S rRNA were selected as reference genes to validate *gdh* gene variation under the conditions of sufficient and insufficient soluble phosphate (Fig. 4). In the absence of soluble phosphate, the *gdh* gene showed the highest level, which was 199.21-fold higher than the gene expression under a soluble phosphate concentration of 10.0 mmol L^-1^. There was no significant difference (P < 0.01) in the *gdh* gene expression level between soluble phosphate concentrations of 5 and 10 mmol L^-1^.

**Fig. 4.**
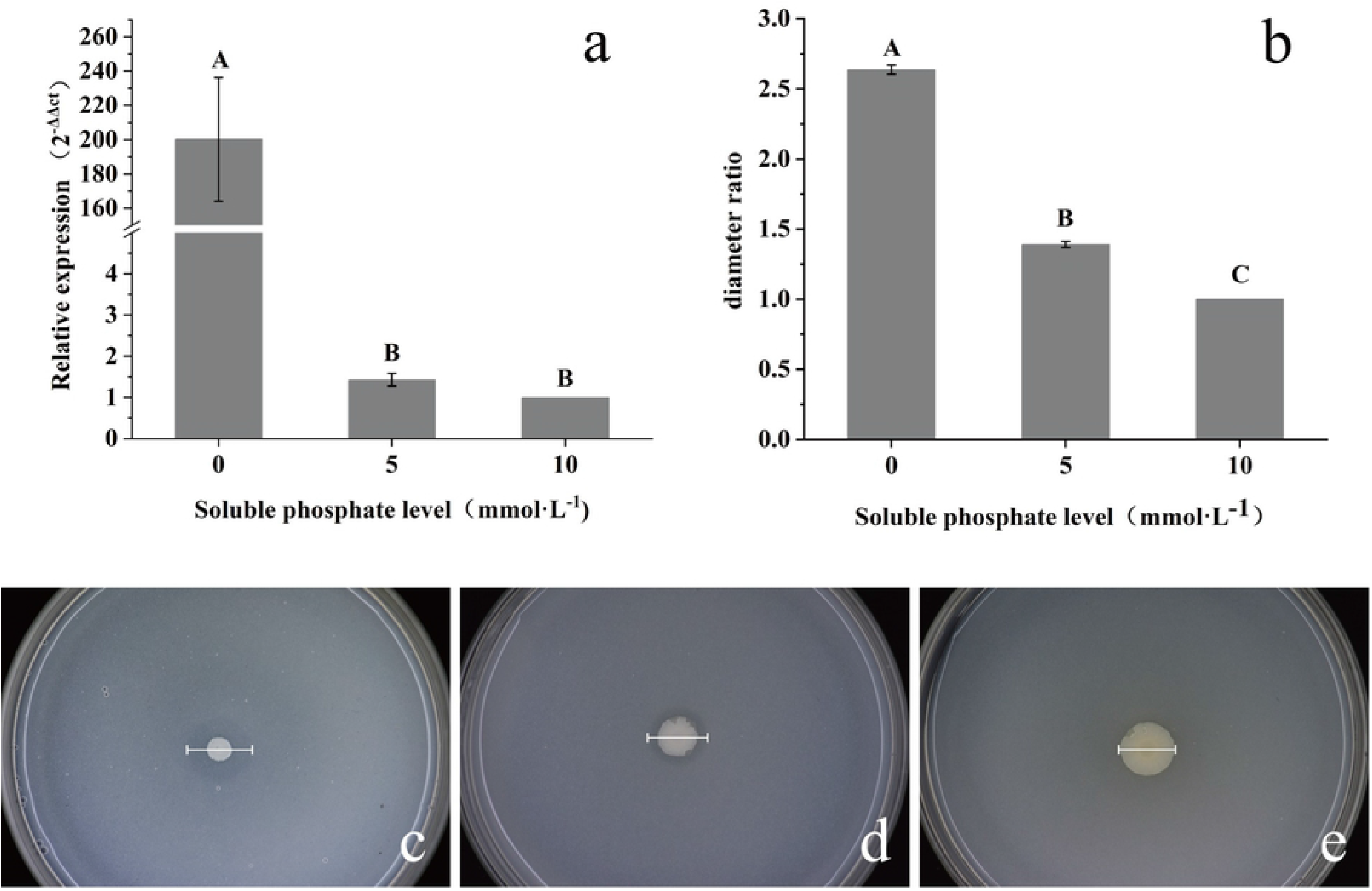
Verification of *gdh* gene expression under different soluble phosphate levels. Relative *gdh* expression levels (a). Colony ratio between the transparent zone diameter and the colony diameter at different phosphate levels (b). Phosphate solubilization on NBRIP plates with different concentrations of soluble phosphate: 0 mmol L^-1^ (c), 5.0 mmol L^-1^ (d), and 10.0 mmol L^-1^ (e).

The phosphate solubilization results on the NBRIP plate are shown in Fig. 4. The phosphate solubilization ability was determined by the ratio between the transparent zone diameter and the colony diameter. At a soluble phosphate concentration of 0 mmol^-1^, the average diameter of the transparent zone was 2.637-fold greater than the diameter of the colony. Compared with the level under soluble phosphate concentrations of 0 and 5 mmol L^-1^, the ratio between the transparent zone diameter and the colony diameter showed a significant decrease (P < 0.01). The ratio decreased with increasing soluble phosphate concentration. No transparent zone was observed at a 10.0 mmol L^-1^ soluble phosphate concentration.

## Discussion

At present, RT–qPCR is used in almost all organisms, such as bacteria [24,26], viruses [21,27], animals [28] and humans [29], and is one of the most widely used methods for gene expression analysis. An increasing number of studies have confirmed that the expression of even the commonly used housekeeping genes may fluctuate in different species, different growing stages and different experimental conditions [27–33]. This suggests that reference genes should be selected according to species and experimental conditions. As the reference genes in the genus *Serratia* have not been reported to date, it is of great significance to confirm the internal reference genes of this genus.

*Serratia ureilytica* is a newly discovered species that was first isolated from the Torsa River in West Bengal, India, in 2005 by Bhaskar [34]. *S. ureilytica* DW2 was recently isolated from the rhizosphere soil of *C. pilosula*. *S. ureilytica* DW2 shows strong phosphorolytic properties and other plant growth-promoting properties, which means that strain DW2 has the potential to be used as a biofertilizer in *C. pilosula* cultivation. When analyzing its phosphate-solubilizing mechanisms based on whole genome sequencing data, it was found that there were no related reference genes for this species, which would limit the quantification of target gene expression under different conditions. The 16S rRNA gene is a common housekeeping gene used to verify the expression of functional genes[35, 36], but the Cq level of 16S rRNA may rely on the cell’s nutritional environment [37]. Furthermore, the expression levels of the target genes in cells were relatively low, but the 16S rRNA content was relatively high, resulting in large differences between the expression levels of 16S rRNA and target genes in the RT–qPCR experiment [38]. Therefore, it was necessary to screen more internal reference genes in *S. ureilytica* DW2.

In the present study, 12 candidate reference genes (16S rRNA, *ftsZ, ftsA, mreB, recA, slyD, thiC, zipA, trpR, rpoD, secA*, and *pyrI*) were selected and cloned for investigation based on previous reports [39–41]. Four genes, *trpR, rpoD, pyrI*, and *secA*, were not selected as candidate genes for internal reference due to poor standard curve data and low transcript abundance. However, according to the results of a study reported by Bai, *rpoD* was a stable internal reference gene in *Pseudomonas brassicacearum* GS20 [41]. Therefore, it could be inferred that the internal reference genes exhibited specificity [42].

Eight genes were finally selected for screening and operation under different conditions. Under the different conditions, the most unstable genes were uniformly identified as *thiC* and *recA* by geNorm and NormFinder, which was in accordance with the findings of previous studies [43, 44].

The most stable genes in this experiment were found to be consistent across all treatment conditions. The top three genes according to geNorm were *zipA*, 16S rRNA, and *slyD*, and the top three genes in NormFinder were *slyD, zipA*, and *mreB*. BestKeeper screened only two stable reference genes that met its screening conditions: *zipA* and 16S rRNA. The results of the present study showed that the inclusion of a third gene had roughly the same effect (V = 0.24) as the use of a second gene (V = 0.22). The use of more than two genes resulted in no significant improvement in the normalization factor. Similar results were reported by Guo [29]. Due to the small differences in V values, two pairs of genes were finally selected as internal references. Therefore, geometric average values were used to rank these candidate reference genes, and *zipA* and 16S rRNA were selected as the most suitable reference genes for strain DW2.

After confirming the reliability of the reference genes, the *gdh* gene was found to be related to the phosphate-solubilizing ability of strain DW2. The *zipA* and 16S genes were then used to validate *gdh* expression under three soluble phosphate concentrations. The expression of *gdh* was upregulated under 0 mmol^-1^ soluble phosphate, which could enhance the ability of DW2 to produce gluconic acid, resulting in the transformation of insoluble phosphate into soluble phosphate. *gdh* expression decreased with increasing soluble phosphorus concentration. When the soluble phosphate content reached 5 and 10 mmol^-1^, *gdh* expression significantly decreased (P < 0.01). This result was consistent with the data obtained from the NBRIP plate, which showed that when the soluble phosphate content was 0 mm, the largest transparent zone diameters were produced. Therefore, the expression of *gdh* could be objectively quantified by *zipA* and 16S, and these two genes could be used as reference genes for *S. ureilytica* DW2. This study was the first to identify reference genes and confirm their reliability in *S. ureilytica* DW2. This study may provide a reference for studies on other strains in the genus *Serratia* and provide a basis for research on phosphate solubilization by *S. ureilytica* DW2.

## Supporting information

**S1 Fig. Agarose gel electrophoresis of PCR products from candidate reference genes.**

**S2 Fig. Melting curve analysis of primer specificity for the amplification of genes from *S. ureilytica* DW2.**

## Acknowledgments

The authors thank the reviewers for their valuable comments on our paper. Yuhe Fan, Pengfei Zhang and Heng Xue helped conduct the *S. ureilytica* DW2 culture experiment. We thank them very much for their hard work.

## Notes

**Conflict of Interest Statement**, We declare that we do not have any commercial or associative interest.

**Fundings:** This work was supported by the General program of Natural Science Foundation of China (32071770), the Fundamental Research Program of Shanxi Province (award No. 202103021223380),the Fund for Shanxi “1331 Project” Key Subjects Construction (1331KSC)

### Competing Interest Statement

The authors have declared no competing interest.

